# Bioavailability of Schisandrin B and its effect on 5-Fluorouracil metabolism in a xenograft mouse model of colorectal cancer

**DOI:** 10.1101/2022.11.28.518277

**Authors:** Pui-Kei Lee, Vanessa Anna Co, Yang Yang, Murphy Lam Yim Wan, Hani El-Nezami, Danyue Zhao

## Abstract

Schisandrin B (Sch-B) is a predominant bioactive lignan in the fruit of a traditional Chinese medicinal plant *Schisandra Chinensis* with widely reported anti-cancer properties. Using a xenograft mouse model of colorectal cancer (CRC), we showed potent anti-tumor effects of Sch-B and synergistic effects when co-treated with the chemotherapy drug, fluorouracil (5-FU). To explore the underlying anti-tumor mechanism of Sch-B, we first compared the bioavailability, metabolism and tissue distribution of Sch-B and its metabolites among healthy and tumor-bearing mice. To understand the drug-phytochemical interactions associated with the synergy between Sch-B and 5-FU, we examined their reciprocal influence on drug metabolism, tissue distribution, and multidrug resistance (MDR) gene expression in tumor-bearing mice. Using a targeted metabolomics approach, three Sch-B metabolites and two bioactive 5-FU metabolites were quantified and found to reach tumor tissue. Generally, Sch-B metabolites were present at higher levels in tumor-bearing than healthy mice, whereas 5-FU metabolite accumulation was remarkably higher in the co-treatment than 5-FU alone group. Moreover, MDR genes were significantly downregulated upon co-treatment, demonstrating the capacity of Sch-B to reverse MDR in chemotherapy. This study showed that Sch-B may serve as a promising adjuvant to chemotherapy drugs via favorably modulating drug metabolism and bioavailability, and attenuating MDR.

## 1. Introduction

*Schisandra Chinensis* is a medicinal plant which has been used as a traditional Chinese medicine (TCM) for thousands of years. Chemical profiling of its herbal extract revealed that the major bioactive components are lignans, a group of polyphenols with two C_6_C_3_ units (Hu et al., 2013). Among them, Schisandrin B (Sch B) is one of the most abundant lignans present in the fruit of *S. Chinensis* (Liu et al., 2013) (**Supplementary Figure 1**). It has been found to exhibit multifarious biological activities, including neuroprotective, cardioprotective, antioxidant, hepatoprotective, anti-inflammatory and anti-cancer effects (Nasser et al., 2020). In particular, Sch B exhibits *in vitro* anti-proliferation and anti-metastasis effects, and *in vivo* anti-tumor activities against various cancers, such as liver, breast, gastric and cervical cancers (Dai et al., 2018; He et al., 2022; Yan et al., 2020). However, to date, little is known about the effects of Sch B on colorectal cancer (CRC).

Colorectal cancer, the third most prevalent cancer, causes approximately 0.94 million deaths worldwide in 2020 (Sung et al., 2021). In terms of CRC treatment, 5-fluorouracil (5-FU) is one of the first-line chemotherapy drugs which mainly targets cancer cells by inhibiting DNA synthesis and RNA functions (Miura et al., 2010). Despite its efficacy and wide application, the administration of 5-FU is often associated with severe side effects, such as diarrhea, vomiting, and stomatitis (Lazar et al., 2004). Hence, development of an alternative therapeutic strategy with reduced side effects is highly demanded. Recently, Sch B has been reported to generate synergy with other chemotherapy drugs with attenuated side effects in gastric and cervical cancers (He et al., 2022; Yan et al., 2020). Our preliminary study showed that oral administration of Sch B in combination with 5-FU significantly reduced tumor volume in a xenograft CRC mouse model with significantly reduced diarrhea symptoms. As such, favorable interactions between Sch B and 5-FU can be expected which may account for the enhanced anti-tumor efficacy and the attenuated side effects in chemotherapy.

Following oral administration of a drug molecule, it will undergo the absorption, distribution, metabolism and excretion (ADME) process which largely decides whether the bioactive molecule can reach the target site(s) and its accumulation levels. For chemotherapy, it is of great importance that the drug can reach and selectively accumulate in the target tumor rather than widely distributed in other tissues. Although earlier studies have investigated the metabolism and bioavailability of Sch B in healthy animals (Wang et al., 2018; Zhu et al., 2013), no study has been conducted in diseased animal models. To understand the mechanism underlying the anti-tumor activity of Sch B and its interaction with 5-FU, it is also imperative to examine its bioavailability and metabolism in tumor-bearing animals. In this study, we aimed to address two key questions to provide insights into the anti-tumor properties of Sch B and the synergy between Sch B and 5-FU. First, are there significant differences regarding the bioavailability, metabolism and tissue distribution of Sch B between healthy and CRC mice? Second, how do Sch B and 5-FU interact with each other which could ultimately lead to altered anti-tumor efficacy? To the best of our knowledge, this is the first study to investigate the bioavailability and metabolism of Sch B and its interaction with the chemotherapy drug 5-FU in a mouse model of CRC.

## 2. Materials and Method

### 2.1 Chemicals and Reagents

All chemicals were of the highest purity available, excepted otherwise specified. Solvents and acids including methanol (Duksan, Ansansi, Korea), acetonitrile (ACN), formic acid (FA) (Macklin, Shanghai, China) and ethyl acetate (Anaqua, Cleveland, Ohio, USA) were all of HPLC grade. β-glucuronidase and fluorouracil (5-FU) were purchased from Sigma-Aldrich (St Louis, Missouri, USA). Standards including schisandrin B (Sch B), schisandrol B (Sch-ol B) and schisantherin A (Sch A) were purchased from Alfa Biotechnology (Chengdu, China), hippuric acid (HA) and ethyl gallate was from Aladdin (Shanghai, China).

### 2.2 Cell culture

The human colorectal cancer cell line HCT-116 (ATCC, CCL-247) was cultured in Dulbecco′s modified eagle′s medium (DMEM) supplemented with 10% fetal bovine serum (FBS, Gibco BRL), and maintained at 37°C under a humidified atmosphere with 5% CO_2_.

### 2.3 Animals

Five-week-old male BALB/c nude mice (Laboratory Animal Unit, The University of Hong Kong) were housed individually with 12 hours light/dark cycle at 22 ± 2 °C. Animals were given *ad libitum* access to a standard diet (AIN-93G, Research Diets, USA) and distilled water. After one-week acclimatization, the mice were divided into CRC and healthy groups. For the CRC mice, 100 μL HCT116 colorectal cancer cells suspension in PBS (10^7^ cell/mL) was mixed with 100 μL Matrigel (Corning, Somerville, Massachusetts USA) and injected subcutaneously to the right flank of the mice and allowed one week for tumor formation. The mice were then randomly assigned to 4 groups, including control (n = 9), Sch B (50 mg/kg) (n = 18), 5-FU (75 mg/kg) (n = 12) and Sch B (50 mg/kg) + 5-FU (75 mg/kg) (n = 12). Sch B was administered by oral gavage every other day, while 5-FU was given by intraperitoneal injection once a week. The control group was administered with phosphate buffered saline (PBS). For the healthy group, animals were treated with 50 mg/kg Sch B by oral gavage every other day (n = 12). The treatment lasted for 14 days for both CRC and healthy groups. All study protocols were approved by the Department of Health, Hong Kong (Ref No.: (19-233) in DH/SHS/8/2/3 Pt.30) and Committee on the Use of Live Animals in Teaching and Research (CULATR no. 5068-19) of the University of Hong Kong.

### 2.4 Biological sample collection

After the 14-day treatment, animals were sacrificed by i.p. injection of sodium pentobarbitone. Plasma and tissue samples (tumor, colon and cecal content) were collected at 6 h following the last dose of Sch B for tumor-bearing mice of all treatment groups (n = 9 for control group and n = 12 for Sch B, 5-FU and Sch B + 5-FU groups) and the healthy group (n = 6). An additional time point at 2 h post-gavage for plasma and tissue samples (tumor, colon and cecal content) collection was assigned for tumor-bearing (n = 6) and healthy groups (n = 6) treated with Sch B alone. Blood was collected from the inferior vena cava in Greiner MiniCollect EDTA tubes (Greiner Bio-one, Kremsmünster, Austria). Plasma was isolated by centrifuging at 2000 x g for 10 min at 4°C, followed by immediate acidification with 2% FA to a final concentration of 0.2%. Tumor and colon tissues were homogenized with or without 0.2% FA at 1:2 (w/v) whereas cecal content was homogenized with water at 1:3 (w/v). All samples were stored at -80°C until analysis.

### 2.5 Extraction of Sch B and metabolites in biological samples

All samples were thawed on ice and extractions were performed at room temperature. Two internal standards (ISs), ethyl gallate (EG) and schisantherin A (Sch A) were diluted in 0.4 M NaH_2_PO_4_ buffer (pH 5.4) and then added to an aliquot of plasma (100 μL), tumor homogenate (200 μL), colon tissue homogenate (200 μL) and cecal content homogenate (500 μL), at a final concentration of 100 ng/mL. The extraction of Sch B and metabolites was performed as per the method of Zhao et al. (2020) with modifications. Briefly, the samples were mixed with 250 μL (for plasma) or 500 μL (for tumor, colon or cecal content) of NaH_2_PO_4_ buffer, followed by digestion with 50 μL (for plasma) or 100 μL (for tumor and colon) of β-glucuronidase (2500 U, in contamination with sulfatase) solution at 37°C for 45 min. Ethyl acetate (500 μL) was added, sonicated in ice-cold water for 5 min and centrifuged at 12000 x g for 5 min. The upper organic phase was collected, followed by adding 20 μL of 2% ascorbic acid in methanol. Two more extractions were performed before the supernatants were pooled and dried under nitrogen. The residue was reconstituted in 100 μL of 80% methanol containing 0.1% FA and sonicated for 5 min. The mixture was then centrifuged at 17,000 x g for 10 min before analyzed by ultra-high-performance liquid chromatography coupled with triple quadrupole mass spectrometry (UHPLC-QqQ/MS).

### 2.6 Instrumentation and analytical methods

The analysis of Sch B, 5-FU and their related metabolites was carried out on an Agilent 6460 triple quadrupole mass spectrometer (Agilent Technology, Palo Alto, CA, USA) with a Water Acquity BEH C18 column (2.1 × 50 mm, 1.7 μm) equipped with a Waters VanGuard Acquity C18 guard column (2.1 × 5 mm, 1.7 μm) (Milford, MA, USA). The mobile phase consisted of 0.1% (v/v) FA in deionized water (A) and 0.1% (v/v) FA in ACN (B). The gradient elution program was set as 0 min, 99% A; 0.5 min, 99% A; 1.5 min, 75% A; 5.5 min, 15% A; 6.8 min, 15% A; 7 min, 96% A. A 3-min post-wash period was included between injections. The flow rate and column temperature were set at 0.45 mL/min and 40 °C, respectively. Each sample was injected by an autosampler set at 4 °C with an injection volume of 3.5 μL.

For MS analysis, the mass spectral data acquisition was obtained under the dynamic multiple reaction monitoring (dMRM) mode with switching polarities. The ESI parameters were as follows: capillary voltage + 3.0 kV/-2.5 kV, nozzle voltage +1.5 kV/-1.0 kV, drying gas temperature 250°C at 12 L/min, sheath gas temperature 250°C at 8 L/min, and a nebulizer pressure at 30 psi. The analytes were identified by comparing the retention time and precursor-product ion transitions with those of the standards. The MS/MS parameters of target metabolites and internal standards were listed in **Supplementary Table 3**. Quantification was achieved with the calibration curves established with the peak area ratio of analyte-to-IS of the quantifiers. All sample extracts were injected into UHPLC-QqQ/MS in a random order and a quality control sample containing all analytes and ISs at a concentration of 100 ng/mL was injected every 10 samples to monitor instrumental stability during sequential runs.

### 2.7 Method Validation

The bioanalysis method was validated in terms of linear dynamic range, lower limit of detection (LLOD), lower limit of quantification (LLOQ), recovery and matrix effect. Calibration curves were obtained by linear plots of peak area ratio (analytes/internal standard) against varying concentrations of each analyte, and linearity (indicated by R^2^) and the linear dynamic range were determined. The LLOD and LLOQ of each analyte were determined as the lowest concentration reaching the signal-to-noise ratio (S/N) ≥ 3 and 10, respectively. Blank plasma and homogenized liver samples from non-treated nude mice were used for assessing recovery and matrix effect. The analytes were spiked at three concentration levels (25, 100 and 400 ng/mL) with two ISs (EG and Sch A) spiked at 100 ng/mL. The recovery (%) was calculated by comparing the mean peak area of analyte spiked pre-extraction and post-extraction with reference to IS peak area. The matrix effect (%) was determined by comparing the peak area of analyte spiked in blank samples post-extraction with that in the reconstituting solvent.

### 2.8 RNA Extraction and RT-qPCR

Total RNA was isolated from tumor using TRI reagent (MRCgene, Cincinnati, OH, USA) according to the manufacturer’s instructions. RNA concentrations and quality were measured using the Thermo Scientific NanoDrop UV-Vis Spectrophotometer (Waltham, MA, USA) and reversely transcribed to cDNA using HiScript™ III RT SuperMix for qPCR (Vazyme Biotech Co., Ltd., Nanjing, China). The mRNA expression levels of multidrug resistance (MDR) genes, including MDR1 and MRP1, were quantified by qPCR with ChamQ SYBR Color qPCR Master Mix (Vazyme Biotech) according to the manufacturer’s instructions. *Beta*-actin was used as the housekeeping gene control. The primer sequences were set with reference to Sun, Xu, Lu, Pan, and Hu (2007). All samples were run on the QuantStudio™ 5 Real-Time PCR System (Applied Biosystems Foster City, CA) in duplicate.

### 2.9 Statistical analysis

Data were expressed as mean ± standard deviation (SD). Normality of data was first assessed by Shapiro–Wilk test. The statistical difference of the data was evaluated by independent t-test, one-way or two-way ANOVA using GraphPad Prism (Version 9.0.0, GraphPad Software, Inc., San Diego, CA). Tukey’s honest significant difference test was used to perform *post-hoc* pairwise comparison between treatment groups. The statistical significance was set at *p* < 0.05.

## 3. Results

### 3.1 Anti-tumor efficacy of Sch B alone or in combination with 5-FU

To evaluate the anti-tumor effects of Sch B alone or in combination with 5-FU, tumor volume was measured at the end of the 14-day treatment **(Supplementary Table 1)**. Mean tumor volume of all three treatment groups was significantly smaller than that of the control group (*p* < 0.001). In addition, although individual treatment with 5-FU resulted in significantly smaller tumor volume than Sch B treatment (*p* < 0.05), we observed an obvious further reduction in tumor volume in the co-treatment group (Sch B + 5-FU) compared with the individual treatment groups (Sch B or 5-FU) **(Figure 1)**.

**Figure 1.**
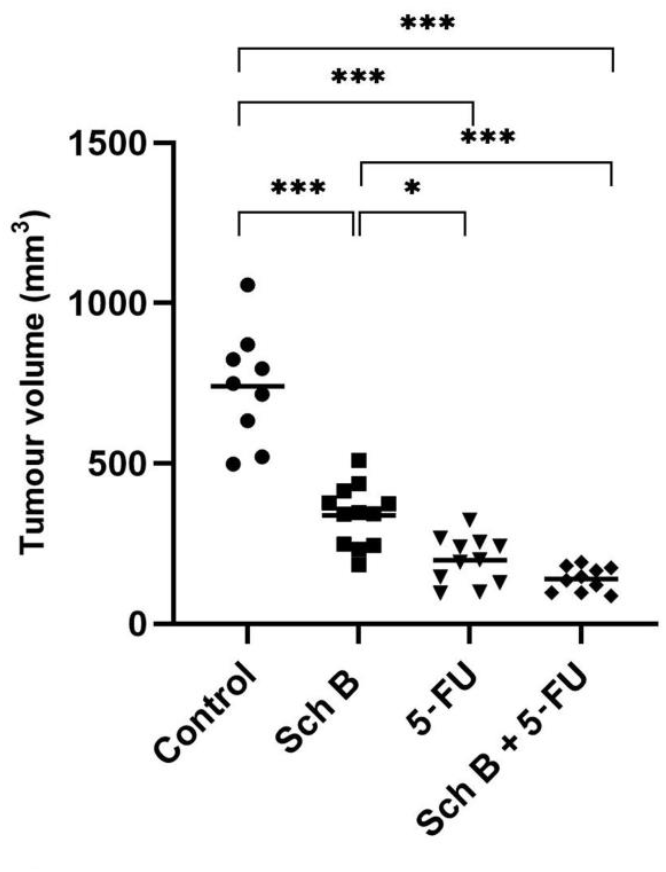
Tumor volume and representative images of tumor of mice with Sch B or 5-FU treatment alone (n = 12) or in combination (n = 12). * *p* ≤ 0.05, ** *p* ≤ 0.01, *** *p* ≤ 0.001 (one-way ANOVA with Tukey’s *post-hoc* test).

### 3.2 Bioanalytical method validation

To confirm the reliability of the newly established UHPLC-QqQ-MS/MS method for quantification of Sch B and its associated metabolites, we validated the bioanalytical method for its calibration linearity, LOD/LOQ, recovery and matrix effects. The validation results are summarized in **Supplementary Table 2**. The calibration curves of all targeted compounds showed satisfactory linearity with a correlation coefficient larger than 0.99. The LLOD and LLOQ of target analytes ranged from 1.1 – 1.2 ng/mL and 2.8 – 3.0 ng/mL, respectively, showing a high sensitivity. The mean recoveries and matrix effects at the three spiking concentrations (25, 100, 400 ng/mL) in blank mouse samples were within 87.2 – 103.6%, indicating satisfactory recoveries and insignificant matrix effects. Based on the validation results, our method has met the acceptance criteria for bioanalysis (FDA, 2018).

### 3.3 Bioavailability and tissue distribution of Sch B in healthy and tumor-bearing mice

Using the validated analytical method, we determined the concentrations of Sch B in plasma, cecal content, colon tissue, and tumor in tumor-bearing mice at 2 h and 6 h after the final oral dose of Sch B (**Figure 2)**. Overall, the levels of Sch B in plasma (47.7 – 330 ng/mL) were significantly higher at 6 h than 2 h post-gavage (*p* < 0.001). For the healthy group, plasma concentration of Sch B at 6 h post-gavage was around 6.3 folds of that measured at 2 h. Similarly, for the CRC group, it was ca. 6.7 folds of that detected at 2 h. However, no significant difference in plasma Sch B concentrations was found between healthy and tumor-bearing mice at both time points.

**Figure 2.**
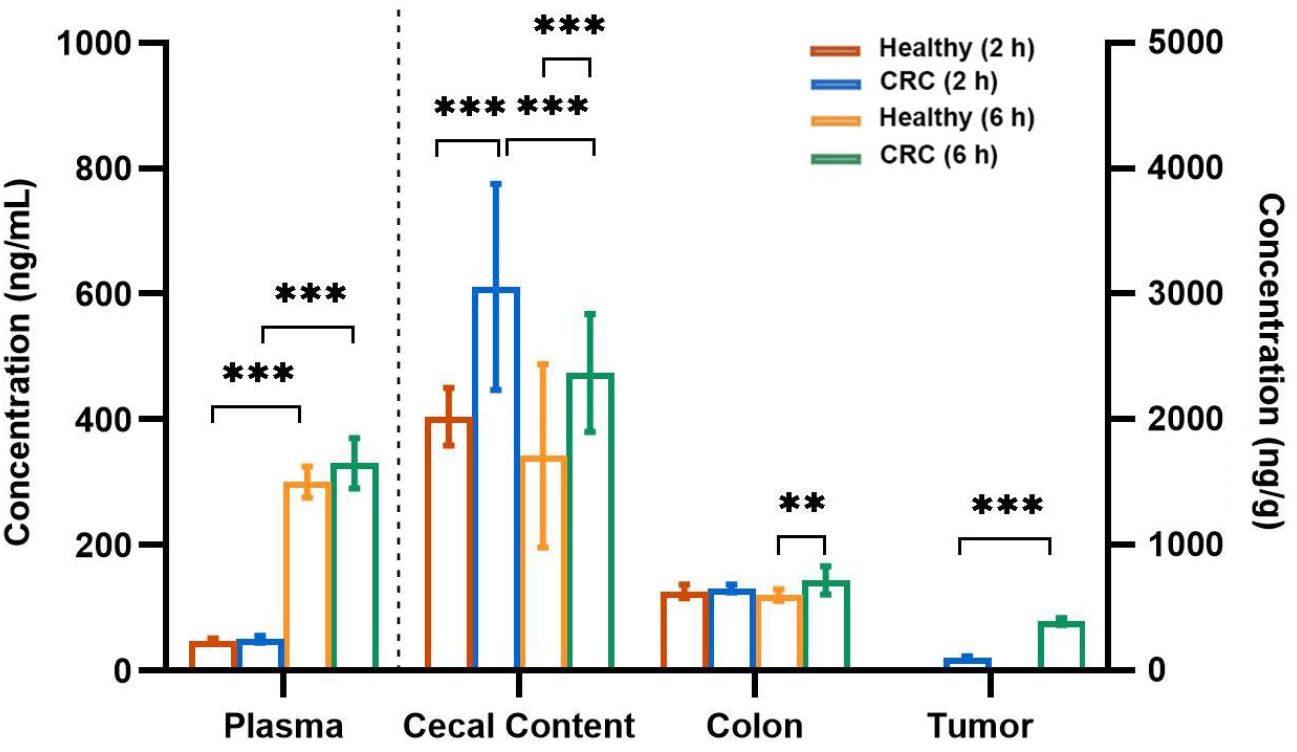
Bioavailability and tissue distribution of Sch B in healthy versus tumor-bearing mice at 2 h and 6 h following the last oral dose of Sch B. Bar graph represents mean ± SD with significant difference evaluated by two-way ANOVA with Tukey’s *post-hoc* test, ** *p* ≤ 0.01, *** *p* ≤ 0.001. Left axis indicates bioavailability of Sch B in plasma (ng/mL) while the right axis indicates tissue distribution of Sch B in cecal, colon and tumor tissue (ng/g).

In addition to the circulating levels, the accumulation of Sch B in cecal content, colon tissue and tumor were also investigated. The average levels of Sch B detected in cecal content, colon tissue, and tumor were within the ranges of 1710 – 3054 ng/g, 601 – 720 ng/g, and 104 – 392 ng/g, respectively. For the healthy group, the Sch B levels in both cecal content and colon tissues were similar between 2 h and 6 h post-gavage. For the CRC mice, the Sch B levels in cecal content at 6 h were remarkably lower (by 28.8%) compared to that of 2 h, while its concentrations in the colon tissue were similar at both time points measured. For the expected target site, i.e., tumor tissue, a significant rise (ca. 4 folds) was observed at 6 h from that of 2 h following Sch B oral administration. When compared between the healthy and CRC groups, a significant difference (ca. 1.5 folds) between the two groups was only observed in cecal content at 6 h post-gavage whereas the Sch B level in colon showed no significant difference.

### 3.4 Metabolism of Sch B in healthy mice and tumor-bearing mice

To compare the metabolic pattern of Sch B between healthy and tumor-bearing mice, the potential metabolites of Sch B upon oral administration in plasma, colon, cecal content and tumor were analyzed. A total of seven Sch B-related metabolites were surveyed, among which four metabolites were detected. Sch-ol B, Gomisin L2 and Gomisin J isomer were regarded as the Sch B-associated metabolites (Qian et al., 2015) **(Supplementary Figure 1)**, and hippuric acid (HA) is a marker metabolite following polyphenol intake (Wishart et al., 2017). Of note, all of them were detected in all sample types, except Gomisin L2 which was not detected in tumor (**Figure 3**).

**Figure 3.**
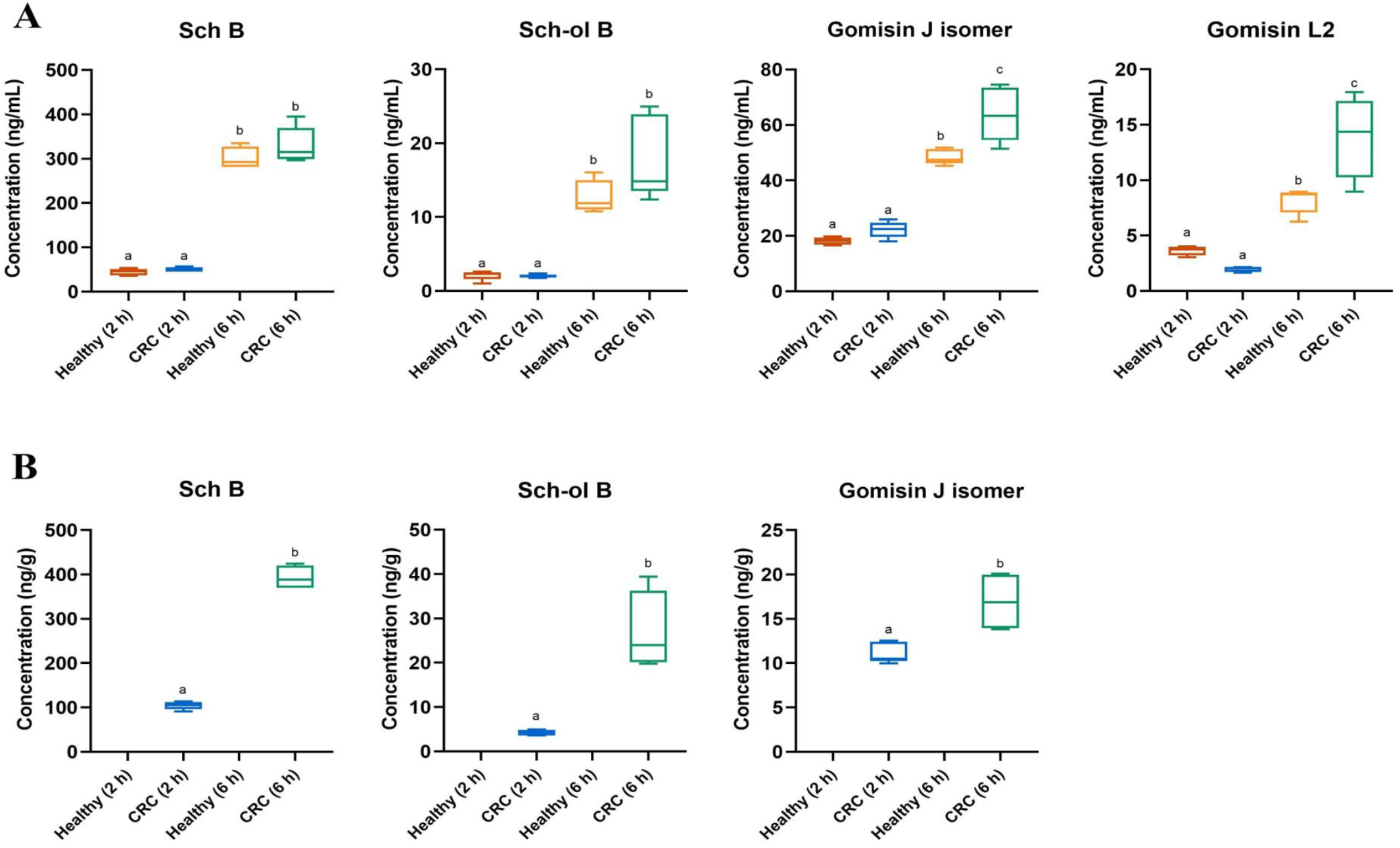
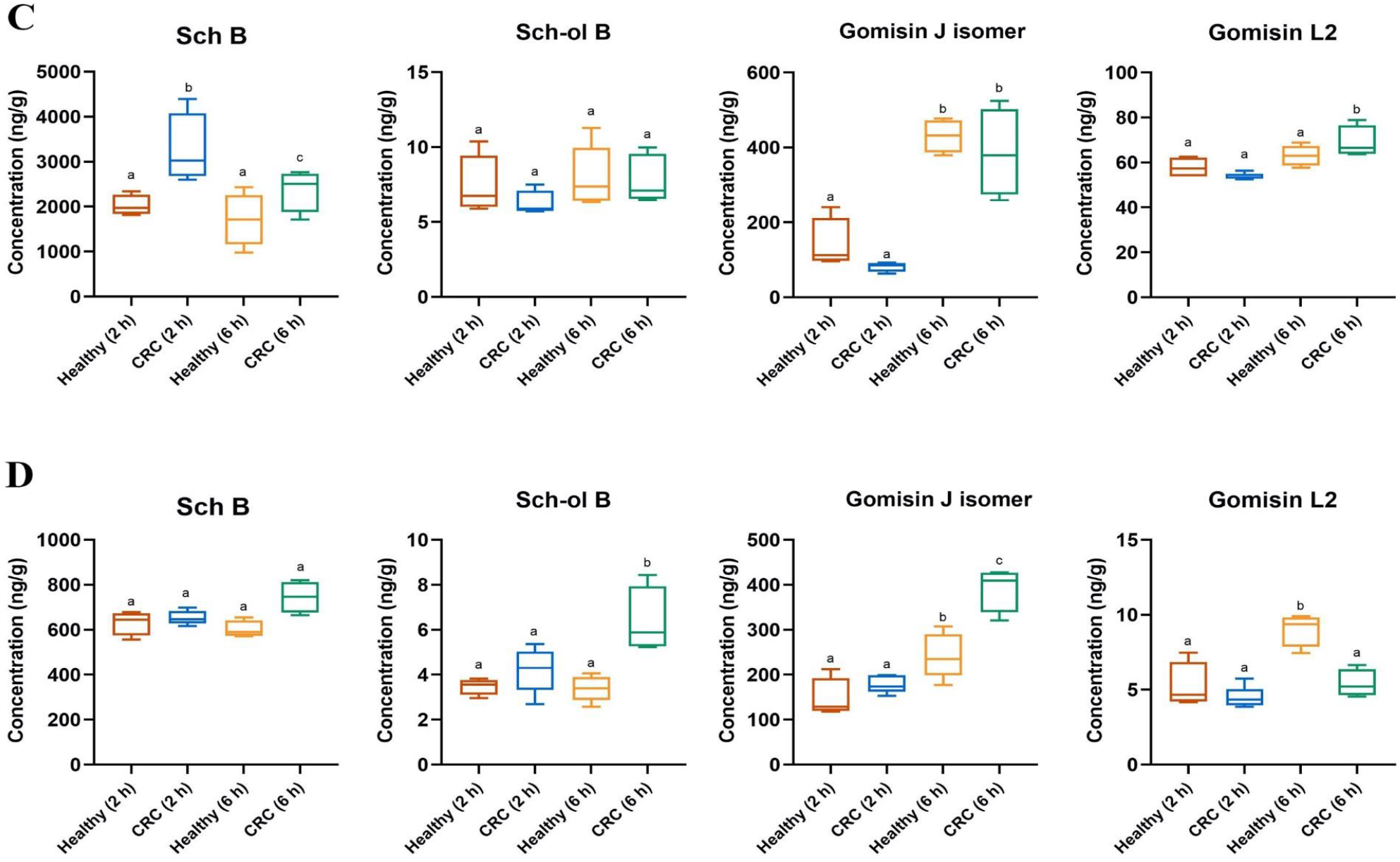
Metabolites of Sch B in (A) plasma, (B) tumor tissue, (C) cecal content and (D) colon tissue of healthy mice and CRC mice at 2 h and 6 h following the last oral dose of Sch B. Different letters represent significant differences in analyte concentrations (*p* < 0.05) between time points and groups; two-way ANOVA with Tukey’s *post-hoc* test, and independent t-test for tumor samples. Sch B, Schisandrin B; Sch-ol B, Schisandrol B.

When comparing between the two time points, for the healthy group, all Sch B metabolites detected in plasma and tumor were present at significantly higher levels at 6 h than 2 h post-gavage (**Figure 3A & 3B)**. Gomisin J isomers levels in cecal and colon tissues were also remarkably higher (ca. 3.0 folds and ca. 1.7 folds respectively) at 6 h than 2 h after the last oral dose of Sch B, while cecal concentrations of Gomisin L2 and Sch-ol B were similar between the two time points respectively (**Figure 3C & 3D)**. For the CRC group, similarly, all Sch B metabolites were mostly found at much higher levels in plasma and tumor samples at 6 h than 2 h post-gavage (**Figure 3A & 3B)**. For Gomisin L2, its cecal content level at 6 h was around 1.5 folds of that measured at 2 h while its levels in colon tissue between the two time points were similar (**Figure 3C & 3D)**. In addition, Sch-ol B level in colon at 6 h post-gavage was also much higher (ca. 1.5 folds) than that at 2 h. Yet, there was no significant difference in its cecal content level between the two time points (**Figure 3C & 3D)**.

We also compared metabolites levels between the healthy and CRC group. The concentrations of the surveyed metabolites in plasma and colon tissue appeared to be similar among healthy and CRC groups at 2 h post-gavage. Interestingly, at 6 h, plasma and colon levels of Gomisin J isomer were significantly higher by 1.3 folds and 1.6 folds, respectively, in CRC group, while those of Sch-ol B showed no difference between the two groups. Gomisin L2 level in CRC group was higher (ca. 2 folds) in the plasma of healthy mice at 6 h post-gavage but was only 60% that in colon tissue. In cecal content, no significant difference in Sch B-associated metabolites was observed between the healthy and CRC groups.

### 3.5 Influence of Sch B administration on 5-FU bioavailability and metabolism in tumor-bearing mice

In view of the reciprocal interactions between Sch B and 5-FU in inhibiting tumor development, we also measured the metabolites levels of 5-FU in plasma, cecal content, colon tissue and tumor, and compared among animals receiving individual or combined treatment (**Table 1**). The native drug molecule, 5-FU, was not detected in any groups at both time points. In addition, two well-referenced active metabolites of 5-FU, namely fluorodeoxyuridine triphosphate (FdUTP) and dihydrofluorouracil (FUH2) (Blondy et al., 2020), were detected in plasma, tumor and colon tissues. In particular, FdUTP, was only detected in the Sch B + 5-FU group, but not in the 5-FU alone group in all types of samples, revealing the influence of Sch B on 5-FU metabolism and bioavailability.

**Table 1.**
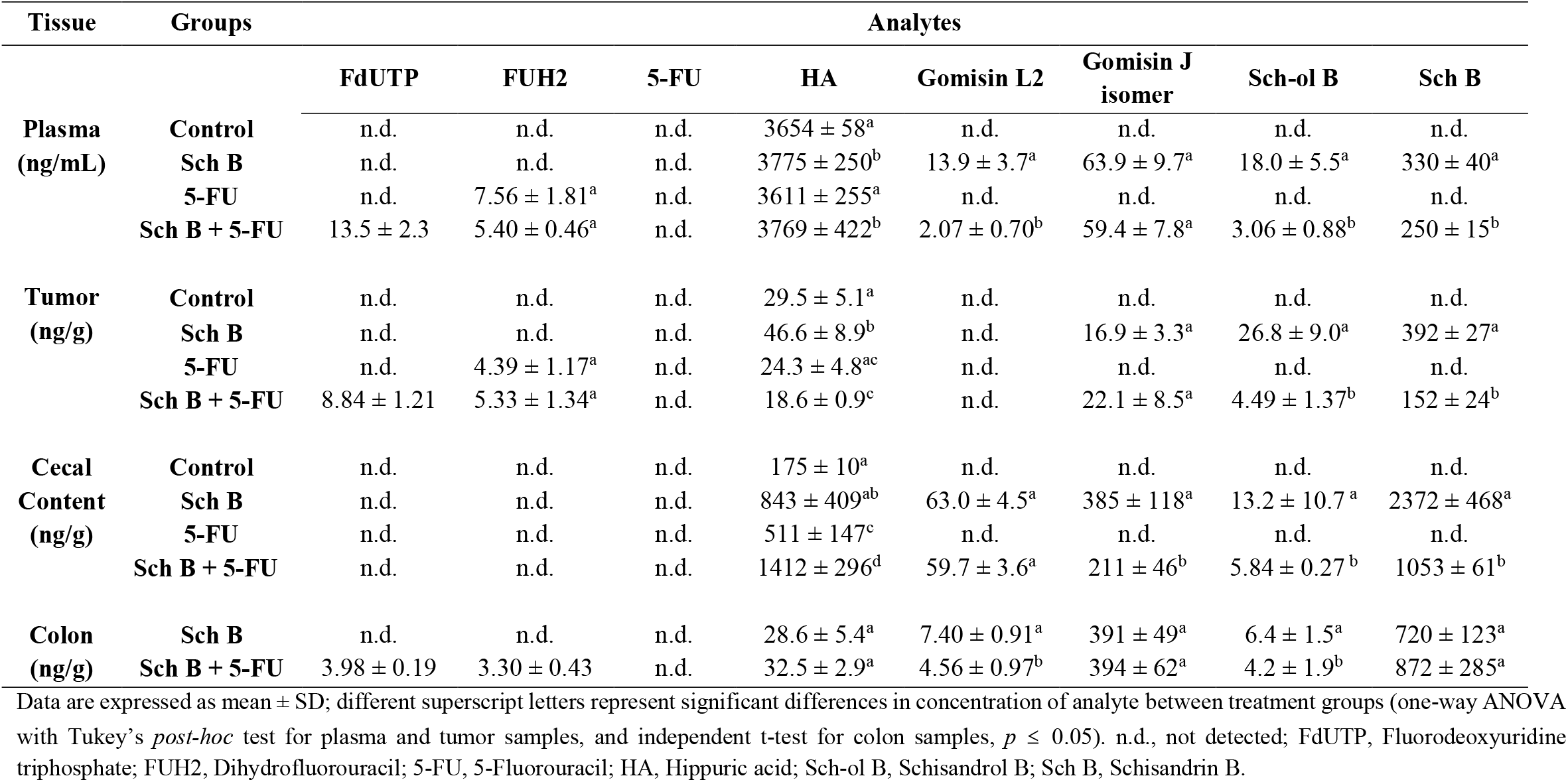
Concentrations of Sch B and 5-FU metabolites in mouse plasma, tumor, cecal content and colon tissues at 6 h following the last dose.

For the two groups treated with Sch B, the concentrations of all Sch B-associated metabolites were lower in the Sch B + 5-FU group than the Sch B alone group, except for Gomisin J isomer whose tumor levels were higher in the Sch B + 5-FU group. This in like manner reveals the influence of 5-FU on Sch B metabolism and bioavailability.

### 3.6 Influence of Sch B administration on the expression of multidrug resistance genes in tumor-bearing mice

To further investigate the anti-tumor action of Sch B and the drug-phytochemical interactions, we explored the effect of Sch B intervention on the expression of key MDR genes in tumor, including MDR1 and MRP1, which were two of the most well-known MDR genes in cancer (Gottesman et al., 2002). The relative mRNA expression of MDR1 and MRP1 is shown in **Figure 4**. As compared to the control, relative expression of MDR1 and MRP1 of the 5-FU group was significantly higher (*p* < 0.01). For the Sch B + 5-FU group, the expression of MDR1 was significantly downregulated compared to the 5-FU group (*p* < 0.01). We also observed a lower expression level of MRP1 in Sch B + 5-FU group when compared to the 5-FU group although the reduction was not significant.

**Figure 4.**
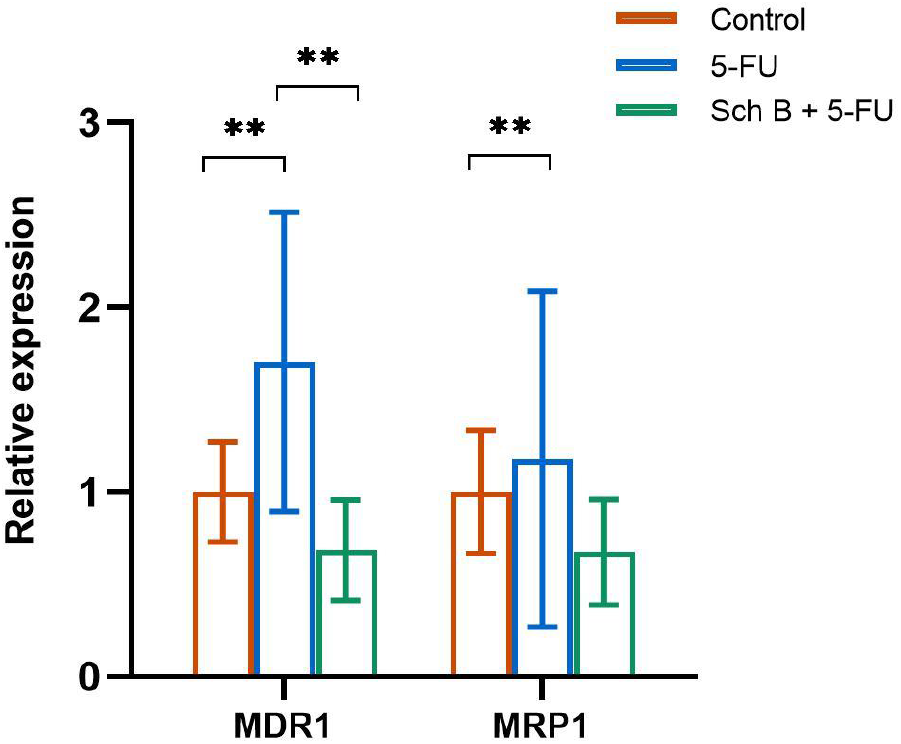
Relative expression of multidrug resistance gene, MDR1 and MRP1, in tumor tissue of Control, 5-FU and Sch B + 5-FU groups. ** *p* ≤ 0.01, one-way ANOVA with Tukey’s *post-hoc* test.

## 4. Discussion

To the best of our knowledge, this is the first study to investigate the bioavailability, metabolism and tissue distribution of Sch B, a major bioactive lignan present in the TCM *Schisandra Chinensis*, in a CRC animal model, and we additionally investigated its reciprocal interactions with 5-FU in attenuating CRC. Our study, using a CRC xenograft nude mouse model, demonstrated potent anti-tumor efficacy of Sch B alone or co-treated with the chemotherapy drug, 5-FU (**Figure 1**). What is more encouraging, when 5-FU was co-administered with Sch B, diarrhea symptoms were remarkably reduced. Our findings suggest the strong potential of Sch B to act as an adjuvant to 5-FU by potentiating anti-tumor efficacy while attenuating side effects in chemotherapy. To provide more insights into the anti-tumor mechanism of Sch B and the synergy between Sch B and 5-FU, we first examined the bioavailability, metabolism and tissue distribution of Sch B in healthy and CRC mice, followed by dissecting the reciprocal interactions between Sch B and 5-FU in CRC mice.

The oral bioavailability and pharmacokinetics behaviors are among the key factors influencing the therapeutic effects of an orally administered xenobiotic compound (e.g., chemotherapy drug-5-FU, or polyphenol-Sch B in our study) (Rein et al., 2013). However, these factors are greatly affected by disease status such as cancer wherein xenobiotic metabolism and excretion are greatly altered (Johnson et al., 2012). Therefore, exploring the difference in xenobiotic metabolism and distribution among individuals under healthy or diseased status could shed light on the anti-tumor mechanisms of Sch B alone and its synergy with 5-FU when applied in combination. Previously, although pharmacokinetic studies of Sch B (as pure compound or medicinal herbal extracts rich in Sch B) were all conducted in healthy rats, large discrepancies in the pharmacokinetic behavior of Sch B were observed. The time for Sch B to reach maximum plasma concentration (T_max_) after oral administration fell within the range of 1-6 h and its half-life (T_1/2_) was around 5-9 h (Wang et al., 2018; Zhu et al., 2013). Based on literature reports, we chose 2 h and 6 h post-gavage as the time points to sacrifice animals and collect samples. In our study, we found significantly higher plasma levels of Sch B at 6 h than at 2 h after oral administration in both healthy and tumor-bearing BALB/c-nude mice (**Figure 2**). These discrepancies may be due to the difference in animal species, genetic makeup and dosing regimen used in the studies compared with previous studies.

It is well known that xenobiotic metabolism can become more extensive under cancerous conditions (Gao & Hu, 2010), and it’s clinically relevant to study the metabolism and tissue distribution of Sch B, the xenobiotic of interest, in a preclinical cancer model. We thus investigated the metabolism of Sch B and tissue distribution of Sch B metabolites in both healthy and tumor-bearing animals. In addition to the tumor, which is the expected target site of action in this study and other cancer-related research, we also examined their accumulation in colon tissue as CRC tumor is normally developed in the colon (**Figure 3**). Among the surveyed metabolites, three Sch B-associated metabolites were detected, including Sch-ol B, Gomisin J and Gomisin L2 (**Supplementary Figure 1**). Hydroxylation in Phase I metabolism was found to be one of the primary biotransformation regimen of *Schisandra* lignans *in vivo* (Wang et al., 2020), during which Sch B was reduced to form Sch-ol B. Of note, Sch-ol B was previously found to show cytotoxicity on HCT116 CRC cells and induced apoptosis by downregulating cyclin D1/cyclin-dependent kinase 4 (CDK4) expression and signaling transduction pathways (Kee et al., 2018). Gomisin L2 and Gomisin J isomer were previously reported as Sch B-associated metabolites *in vitro*, which could be formed following demethylation of Sch B by the gut microbiota (Qian et al., 2015). When comparing between the healthy and CRC groups, both Gomisin L2 and Gomisin J isomer were significantly higher in the plasma and colon tissue of tumor-bearing mice than in healthy mice. The higher levels of Sch B-associated metabolites, particularly the Gomisin isomers, in tumor-bearing mice revealed the influence of cancerous state on Sch B metabolism and bioavailability under the experimental conditions. In addition to Sch B, other metabolites such as Sch-ol B and Gomisin J isomer were also detected in tumor tissue. Previously, dietary lignans, such as secoisolariciresinol diglucoside, were reported to reach tumor tissue to exhibit a direct anti-tumor effect (Saarinen et al., 2008). We are the first to report that Sch B, a TCM-derived bioactive lignan, and its metabolites also accumulate substantially in tumor tissue.

As we noticed obvious drug-phytochemical interactions that may in part account for the synergistic anticancer efficacy, we further explored the interactions from two perspectives, i.e., the bioavailability and metabolism of xenobiotics under the influence of Sch B and/or 5-FU, and the expression of MDR genes. Upon co-treatment with Sch B and 5-FU, we observed significantly lower concentrations of Sch B-associated metabolites in plasma and tissues than those of Sch B treatment alone. Interestingly, Gomisin J was detected in the group treated with Sch B + 5-FU, but not in the Sch B alone group. This suggests unique drug-phytochemical interactions which can contribute to the more notable reduction in tumor size upon the co-treatment. In addition, following oral administration and intestinal absorption, similar to all xenobiotics, the non-absorbed proportion of Sch B reaches the large intestine (cecum and colon), which is one of the major sites of xenobiotic catabolism due to the high abundance of microorganisms present. Hence, we also measured the accumulation of gut-derived bioactive metabolites of lignans, such as enterolactone and enterodiol (Mali et al., 2019), in our samples. We detected enterolactone in all sampled tissues and their levels were notably higher in the group treated with Sch B + 5-FU than those of Sch B alone group. Previously, *in vivo* studies have revealed diminished microbial diversity upon chemotherapy treatments. In particular, 5-FU decreased the abundance of commensal bacteria in the gut leading to dysbiosis (Vanlancker et al., 2017). As gut microbiota homeostasis is crucial to gut health and immune system integrity, co-administration of Sch B could help restore microbiome symbiosis and relieve the severe inflammation induced by the chemotherapy drug (Zheng et al., 2020). To further understand the involvement of the gut microbiota, we are working on *in vitro* anaerobic fermentation studies simulating the microbial metabolism of Sch B in the colon, and metagenomics analysis will also be conducted. We have identified new metabolites while the characterization is still on-going.

Mounting evidence has shown that multidrug resistance is one of the prominent factors limiting chemotherapy efficacy. This is largely due to enhanced xenobiotic metabolism and drug efflux of tumor cells in response to chemotherapy drugs (Wu et al., 2014). Earlier studies have revealed the ability of Sch B to reverse P-glycoprotein and MDR1-mediated drug resistance and efflux *in vitro* (Pan et al., 2005; Sun et al., 2006). To confirm such drug sensitizing property *in vivo*, we examined the expression levels of major MDR genes hoping to further elucidate the drug-phytochemical interaction. Two of the most well-known multidrug resistance proteins in cancer include P-glycoprotein (MDR1 expression) and MDR-associated protein 1 (MRP1 expression) were assessed. In this study, we found remarkable reductions in MDR1 and MRP1 mRNA expression in the tumor tissue of CRC mice co-treated with Sch B and 5-FU (**Figure 4**), which may account for the higher levels of the active metabolites of 5-FU in the co-treatment group (**Table 1**). This can be largely attribute to Sch B’s capacity to reverse the adversely affected ADME processes (Krishna & Mayer, 2000) and facilitate the uptake of the chemotherapy drug. For example, we found that oral administration of Sch B facilitated tumor penetration of FdUTP, an active form of 5-FU transformed by thymidine phosphorylase enzyme. FdUTP was reported to suppress tumor DNA synthesis and induce apoptosis of tumor cells (Blondy et al., 2020). Of note, FdUTP was detected only in Sch B + 5-FU group, but not in 5-FU group. Thus, Sch B could possibly enhance the conversion of 5-FU into FdUTP, leading to accelerated DNA damage and apoptosis of tumor cells. Collectively, we showed the potential of Sch B to attenuate CRC cell resistance to 5-FU and also to extend the exposure period locally (in tumor tissue) and systemically, resulting in the significant reduction in tumor volume.

Despite the new insights into the bioavailability, metabolism, tissue distribution and drug-phytochemical interactions between Sch B and 5-FU in a well-accepted animal model of CRC, there are some limitations of this study. For example, information on Sch B excretion is lacking which is important for assessing the overall bioavailability, and more novel Sch B metabolites are yet to be uncovered. In addition, the location of tumor is in the flank rather than in the colon lumen, an orthotropic CRC model may be needed to confirm the tumor-targeting ability of Sch B. Further studies to identify more phase II and/or microbial-derived metabolites of Sch B are warranted to uncover its metabolic pathways and the bioactive forms of this phytochemical.

## 5. Conclusion

In conclusion, the present study revealed significant anti-tumor efficacy of Sch B alone and in combination with the chemotherapy drug, 5-FU. Using a targeted metabolomic approach, three Sch B–associated metabolites were identified and quantified. We also confirmed the ability of Sch B and its associated metabolites to reach tumor tissue, potentially contributing to the anti-tumor efficacy. Significantly higher levels of Sch B and some metabolites were detected in tumor-bearing compared with healthy mice. In addition, we dissected the drug-phytochemical interactions in terms of xenobiotic metabolism and degree of multidrug resistance. The significantly reduced multidrug resistance gene expression upon co-treatment of Sch B on top of 5-FU could be positively linked to the enhanced chemotherapy efficacy. All in all, this study demonstrated the synergistic anti-tumor effects of Sch B (a TCM-derived phytochemical) and 5-FU (a classical chemotherapy drug) in treating CRC, corroborating the promising potential of integrating Chinese and Western medicines for more pronounced efficacy and lower side effects in cancer therapy.

## Acknowledgement

The authors wish to acknowledge the support from the Department of Applied Biology and Chemical Technology (ABCT) and Research Center for Chinese Medicine Innovation (RCMI) at the Hong Kong Polytechnic University (PolyU). The skillful technical support from Dr. Pui-Kin So from the University Research Facility in Life Science at PolyU is also greatly appreciated. This work was supported by the Exploratory Projects of RCMI (P0041459) and PolyU Start-up Fund (P0032176).

## Data Availability Statement

The data that support the findings of this study are available from the corresponding authors upon reasonable request.

