## Supplementary Table 1-3 for "Bioavailability of Schisandrin B and its effect on 5-Fluorouracil metabolism in a xenograft mouse model of colorectal cancer"

### Supporting Information

**Supplementary Table 1.** Final body weight and tumor volume of BALB/c-nude mice.

| Groups | n | Body weight (g) | Tumor volume ($mm^3$) |
|---|---|---|---|
| ***Tumor-bearing mice*** | | | |
| Control | 9 | $20.1 \pm 1.2^{a}$ | $740 \pm 175^{a}$ |
| Sch B | 18 | $21.4 \pm 1.0^{a}$ | $337 \pm 95^{b}$ |
| 5-FU | 12 | $17.2 \pm 2.7^{b}$ | $198 \pm 74^{c}$ |
| Sch B + 5-FU | 12 | $17.3 \pm 2.2^{b}$ | $139 \pm 38^{c}$ |
| ***Healthy mice*** | 12 | $24.1 \pm 0.9^{c}$ | - |

Data are expressed as mean ± SD. Different superscript letters represent significant differences in body weight or tumor volume between treatment groups (one-way ANOVA with Tukey's *post-hoc* test, $p \leq 0.05$).

**Supplementary Table 2.** Analytical method validation parameters of targeted compounds in blank mouse samples.

| Analytes | Retention time (min) | Calibration Curve | $R^2$ | Dynamic linear range (ng/mL) | LLOD (ng/mL) | LLOQ (ng/mL) | Spiked amount (ng/mL) | Recoveries (mean ± SD, %) | | Matrix effects (mean ± SD, %) | |
|---|---|---|---|---|---|---|---|---|---|---|---|
| | | | | | | | | Plasma | Liver | Plasma | Liver |
| Sch B | 5.72 | y = 24.555x – 0.044 | 0.9985 | 1.1 - 1100 | 1.1 | 2.8 | 25 | 98.9 ± 3.0 | 90.1 ± 1.8 | 93.5 ± 5.6 | 90.0 ± 1.3 |
| | | | | | | | 100 | 102.2 ± 2.7 | 92.1 ± 2.8 | 94.3 ± 5.2 | 96.4 ± 2.2 |
| | | | | | | | 400 | 100.7 ± 1.5 | 93.2 ± 4.5 | 94.2 ± 4.9 | 98.2 ± 1.2 |
| Sch-ol B | 4.32 | y = 9.010x – 0.069 | 0.9977 | 1.1 - 1100 | 1.1 | 2.8 | 25 | 93.7 ± 3.8 | 87.2 ± 1.7 | 93.1 ± 3.7 | 91.8 ± 4.5 |
| | | | | | | | 100 | 91.2 ± 2.1 | 87.6 ± 1.9 | 100.3 ± 0.3 | 91.3 ± 2.2 |
| | | | | | | | 400 | 97.8 ± 3.2 | 90.7 ± 2.2 | 103.6 ± 2.0 | 95.9 ± 4.4 |
| HA | 1.94 | y = 8.771x + 0.056 | 0.9997 | 1.2 - 1200 | 1.2 | 3.0 | 25 | 98.9 ± 3.0 | 100.2 ± 1.8 | 93.5 ± 5.6 | 99.3 ± 3.2 |
| | | | | | | | 100 | 102.2 ± 2.7 | 89.7 ± 3.4 | 94.3 ± 5.2 | 94.4 ± 3.3 |
| | | | | | | | 400 | 100.7 ± 1.5 | 92.5 ± 4.8 | 94.2 ± 4.9 | 92.3 ± 4.8 |

**Abbreviations:** HA, Hippuric acid; Sch B, Schisandrin B; Sch-ol B, Schisandrol B.

**Supplementary Table 3.** Optimized MS/MS parameters of target metabolites surveyed in this study and internal standards.

| **Compounds** | **Molecular weight (g/mol)** | **ESI polarity** | **MRM transitions Precursor → Product ion^ (m/z)** | **Fragmentor voltage (V)** | **Collision energy (V)** |
|---|---|---|---|---|---|
| Sch B | 400.5 | + | 401.2 → 300.1 (285.1) | 160 | 20 |
| Sch-ol B | 416.5 | + | 399.1→ 368.2 (330.0) | 110 | 20 |
| Gomisin L2 | 386.5 | + | 387.2 → 286.1 (324.1) | 160 | 20 |
| Gomisin J isomer | 388.5 | + | 389.2 → 288.1 (374.2) | 160 | 20 |
| Sch B metabolite 1 ($C_{23}H_{28}O_7$) | 416.2 | + | 417.2 → 316.1 (402.2) | 160 | 20 |
| Sch B metabolite 2 ($C_{28}H_{36}O_{12}$) | 564.2 | + | 565.2 → 389.2 (357.2) | 160 | 20 |
| HA | 179.1 | - | 178.1 → 134.1 (77.0) | 90 | 8 |
| 5-FU | 130.0 | - | 129.0 → 42.2 (86.0) | 90 | 15 |
| FdUTP | 486.0 | - | 405.0 → 79.0 (158.9) | 90 | 15 |
| FUH2 | 132.1 | - | 131.1 → 82.9 (41.6) | 90 | 15 |
| FdUMP | 326.0 | - | 325.0 → 79.0 (129.0) | 90 | 15 |
| FUTP | 502.9 | - | 501.9 → 158.9 (273.0) | 90 | 15 |
| FUPA | 150.1 | + | 151.1 → 90.1 (108.1) | 90 | 15 |

| | | | | | |
|---|---|---|---|---|---|
| FBAL | 107.1 | + | 108.0 → 62.1 (90.1) | 90 | 15 |
| Enterolactone | 298.0 | - | 297.0 → 253.1 (106.9) | 140 | 15 |
| Enterodiol | 302.1 | - | 301.1 → 106.0 (253.0) | 140 | 15 |
| EG (IS 1) | 198.2 | - | 197.0 → 124.0 (169.3) | 110 | 25 |
| Sch A (IS 2) | 536.6 | + | 554.2 → 415.2 (371.2) | 90 | 20 |

^ Product ion refers as quantifier ion. Qualifier ion is shown in parentheses.

**Abbreviations:** Sch B, Schisandrin B; Sch-ol B, Schisandrol B; HA, Hippuric acid; 5-FU, 5-Fluorouracil; FdUTP, Fluorodeoxyuridine triphosphate; FUH2, Dihydrofluorouracil. IS (internal standard): EG, Ethyl gallate; Sch A, Schisantherin A.

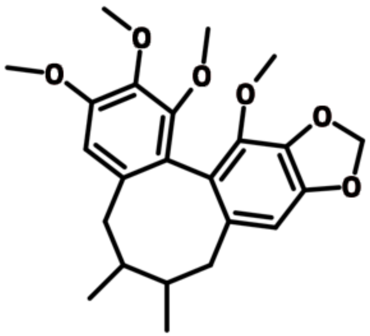

**Schisandrin B (Sch B)**

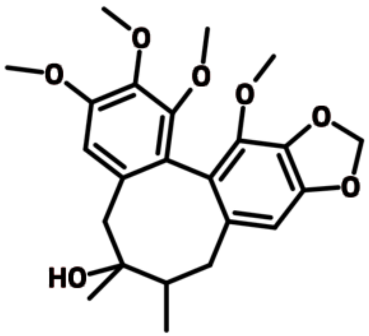

**Schisandrol B (Sch-ol B)**

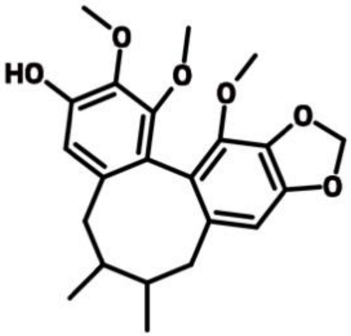

**Gomisin L2**

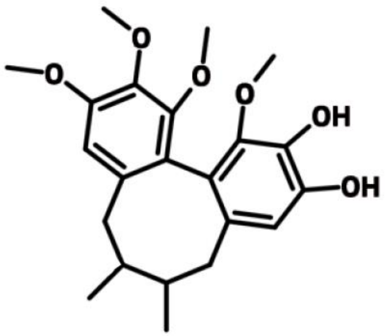

**Gomisin J isomer**

**Supplementary Figure 1**. Molecular structure of Sch B and its associated metabolites detected in this study.
